# Natural language processing models reveal neural dynamics of human conversation

**DOI:** 10.1101/2023.03.10.531095

**Authors:** Jing Cai, Alex E. Hadjinicolaou, Angelique C. Paulk, Daniel J. Soper, Tian Xia, Ziv M. Williams, Sydney S. Cash

## Abstract

Through conversation, humans relay complex information through the alternation of speech production and comprehension. The neural mechanisms that underlie these complementary processes or through which information is precisely conveyed by language, however, remain poorly understood. Here, we used pretrained deep learning natural language processing models in combination with intracranial neuronal recordings to discover neural signals that reliably reflect speech production, comprehension, and their transitions during natural conversation between individuals. Our findings indicate that neural activities that encoded linguistic information were broadly distributed throughout frontotemporal areas across multiple frequency bands. We also find that these activities were specific to the words and sentences being conveyed and that they were dependent on the word’s specific context and order. Finally, we demonstrate that these neural patterns partially overlapped during language production and comprehension and that listener-speaker transitions were associated with specific, time-aligned changes in neural activity. Collectively, our findings reveal a dynamical organization of neural activities that subserve language production and comprehension during natural conversation and harness the use of deep learning models in understanding the neural mechanisms underlying human language.

## Introduction

Our ability to communicate through language requires two distinct but complementary linguistic computations: comprehension and production. Speech comprehension involves a structured succession of processes that extract information from acoustic-phonetic signals allowing us to understand the meaning of sentences and to comprehend the thematic and contextual information being conveyed *(1-6)*. In contrast, speech production planning involves a reverse process whereby higher-order conceptual information is converted to motor planning features for articulation *(7-9)*. Together, these processes are necessary for communicating information during conversations that usually involve rapid alternations between speakers every few seconds *(10, 11)*.

Our primary mode for understanding human language has been through highly structured tasks under artificial constraints that do not capture the dynamics of conversations in a natural scenario *(12)*. Blocked design tasks, in particular, involve predetermined language materials and scripted turn-taking in conversations *(13-15)*. This results in content and timing that are often unnatural. Furthermore, due to the repetition of identical trials, subsequent statistical analyses may become contingent on the number of repetitions, rendering the findings less applicable to natural conversations. Together, these task settings are inherently constrained and insufficient for revealing the neural foundations of complex cognitive activities, particularly those associated with spontaneous and free-flowing speech *(16)*. As a result, how linguistic information is precisely represented in the brain during natural conversation or what common neural processes subserve speech production and comprehension have remained a challenge to understand. Whereas certain brain areas have been shown to distinguish well-formed meaningful sentences from degraded or non-meaningful stimuli, suggesting their involvement in language processing *(17-20)*, the neural process by which linguistic information may be represented in the brain has been difficult to study directly. Further, while there is often broad overlap between areas involved in language production and comprehension *(21-24)*, whether common neural processes are involved in representing linguistic information when speaking and listening during conversation remains poorly understood. Finally, although certain areas of the brain have been implicated in conversation or turn-taking *(13, 25-27)*, little is known about whether or how brain activity relates to information conveyed during dialogue, especially when considered with the temporal dynamics of natural speech since the methods for analyzing natural communication are still limited.

The unrestricted and boundless character of natural conversation poses a significant challenge in comprehensively representing language, thereby complicating the study of neural mechanisms underlying this cognitive process. The recent advancement of natural language processing (NLP) models based on artificial deep learning neural networks has provided a new prospective platform by which to begin studying continuous, natural linguistic interactions. These models have been shown to display high-level performative interactions with human subjects in conversations *(28)*, and can achieve state-of-the-art benchmarks in comprehension-based tasks and question-answering *(29-31)*. Indeed, these models are capable of capturing specific word sequences and their composition within phrases and sentences through hierarchical layers in a way that can be potentially compared to neural activity *(32, 33)*. NLP models have also demonstrated high performance in explaining brain activity during passive listening *(34-36)*, suggesting their neurobiological plausibility. Here, we utilize these models as artificial, hierarchically structured vectorized representations of language during natural dialogue in order to investigate how linguistic information is represented in the brain during conversation.

## Results

### Neural recordings during natural conversation

Local field potential (LFP) recordings were obtained using semi-chronically implanted depth electrodes in 14 participants undergoing epilepsy monitoring as part of their clinical care (6 females and 8 males, average age of 34, ranging between 16 to 59, **Fig. 1a, Table S1**). Together, we recorded from 1910 bipolar-referenced channels (**Fig. 1b**). These channels spanned a total of 39 brain areas across both hemispheres (**Table S2**). Electrodes that displayed low signal-to-noise or frequent epileptiform discharges were excluded (Methods). For all recordings, LFPs were filtered and transformed to envelopes at alpha (8-13 Hz), beta (13-30), low gamma (30-55), mid-gamma (70-110), and high gamma (130-170) frequency bands (**Fig. S1**).

**Fig. 1.**
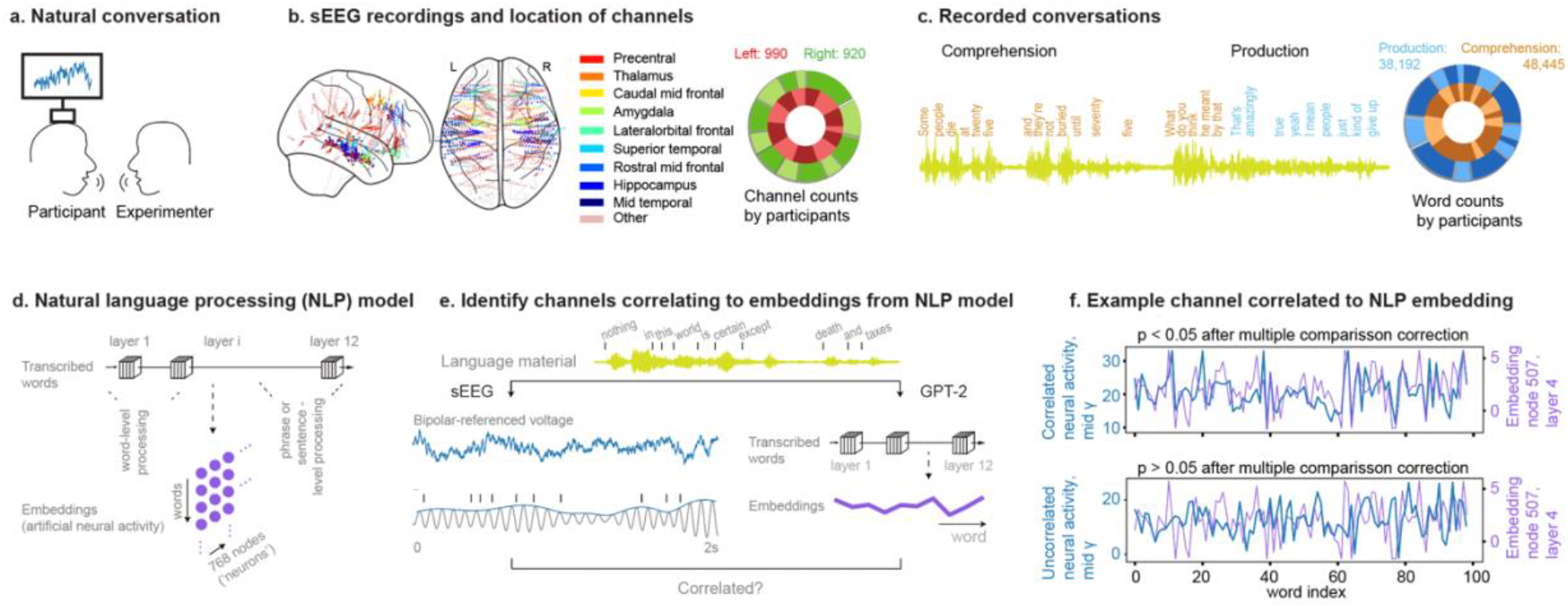
Channel distributions and speech transcription. **a**. Participants were instructed to have natural conversations with an experimenter with their neural activities recorded. **b**. Depth electrodes were implanted in fourteen participants to record local field potentials (LFP). A total of 1910 bipolar-referenced channel channels that demonstrated a low rate of epileptiform discharges were primarily distributed in the frontal, temporal and mesial areas in both hemispheres. The locations of the channels are shown in the *left*, and the number of channels for each participant is shown on the *right* by brightness. **c**. The conversation materials were transcribed to words, and whether a word was perceived or produced by the participant was labelled *(left)*. This resulted in a total of 86,637 words among all participants, and the word counts from each participant is shown on the *right* by brightness. **d**. The transcribed words perceived and produced by the participants were input to the GPT-2 model, and the embeddings from all nodes and all layers were obtained. **e**. These embeddings were correlated to the LFPs at each frequency band. In this way, we identify channels that were significantly correlated to artificial activities of specific nodes in the artificial model. **f**. Example channel that significant correlated to an artificial embedding *(top)* and not correlated to the artificial embedding *(bottom)*.

During recordings, the participants engaged in unconstrained, free-flowing conversation with an experimenter for approximately an hour (range 16 to 92 minutes; Methods). These conversations ranged broadly in topic and theme, allowing individuals to both listen and speak (**Table S3**). All transcribed words were synchronized to neural activities at millisecond resolution. These included 2728 ± 1804 (mean ± s.t.d) words during production and 3460 ± 2581 words during comprehension (**Fig. 1c**). There were an average of 168 transitions between listening and speaking; together reflecting the dynamic interchange between the individuals involved.

### Cerebral network activity patterns in comparison to NLP model-based activity

To quantify the degree to which neural activity reflected the information being conveyed during dialogue (e.g., rather than simply any changes in activity when speaking or listening), we employed a pre-trained GPT-2 (small) model capable of capturing the variance of brain activity in relation to linguistic information being conveyed *(37-39)*. This model also importantly provided access to the hidden embeddings at scales that are tenable for neuronal analysis.

Here, the model embeddings (a set of hierarchically organized vectors serving as the artificial ‘neural’ activities) were trained to represent linguistic features extracted from vast language corpora. When applied to a natural conversation, this model can then vectorize the linguistic information in a quantitative manner which can be directly compared to simultaneously obtained neural data *(34, 40)* (**Fig. 1d, e**). Thus, for example, if the pattern of neural activity in a particular brain area consistently matched the activity derived from an artificial neural activity in these language models, this would mean that their patterns carry linguistic information about the conversations being relayed (see Methods). Specifically, we used the same words that participants spoke or listened to as input to the artificial model and examined correlations between brain and model activity across words to elucidate neural activities that participated in language processing.

By tracking LFP signals across broadly varying conversations, we found that changes in activity in widespread parts of the brain were consistently aligned with those of the NLP model (**Fig. 1d, e, f**). When accounting for all recording contacts and frequency bands, there was an overall wide distribution of correlated channels across most brain areas and frequency bands with the proportion of these channels being significantly higher than expected from chance (Chi-square test, statistics = 7785, p < 10^-100^). The mean and standard deviation of R-value of all channels that displayed significant correlation was 0.12 ± 0.04 for speaking and 0.10 ± 0.03 for listening (**Fig. 2a, b**), meaning that variations neural activity reflected information captured by the model’s embeddings.

**Fig. 2.**
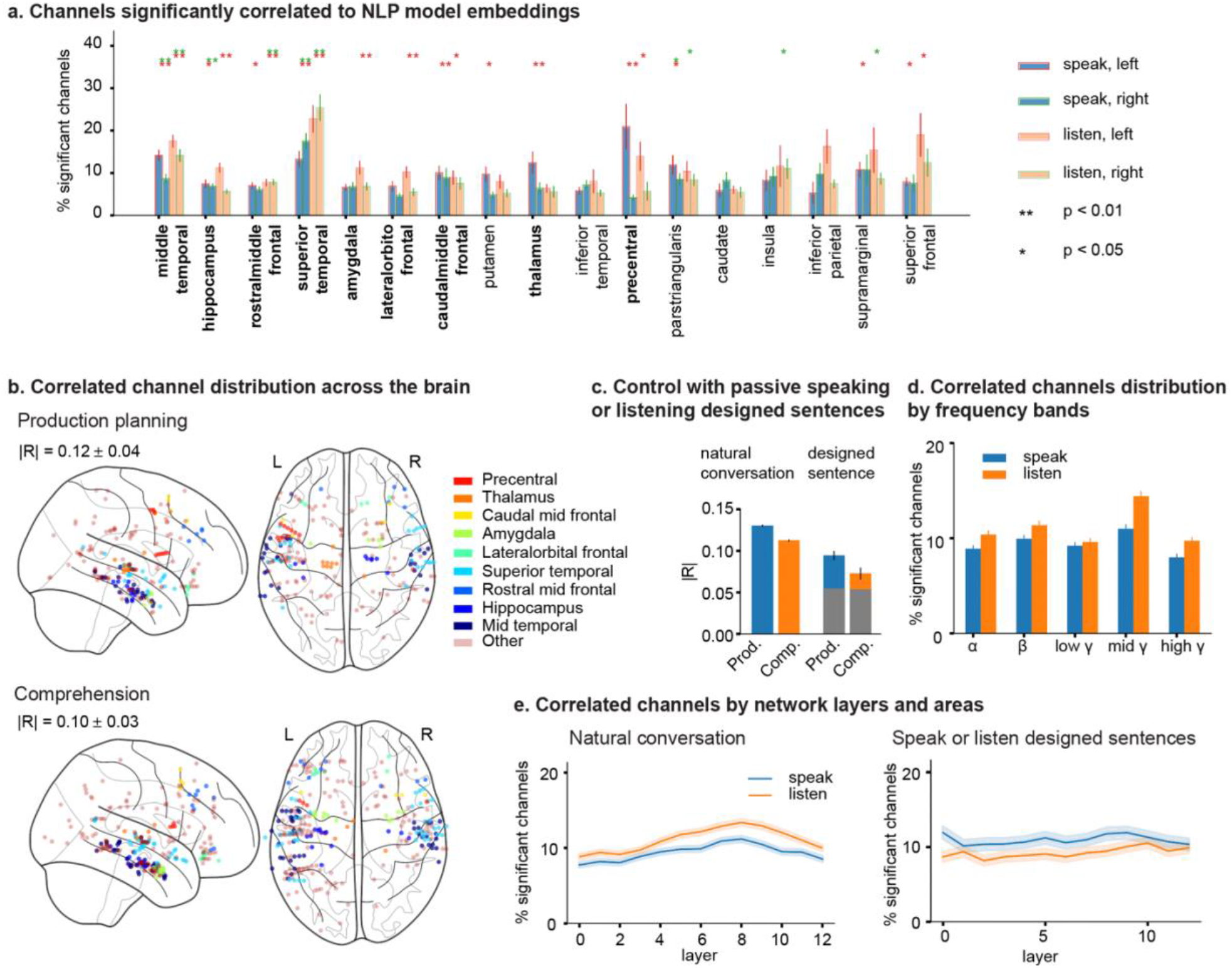
Correlating brain activities to a pre-trained natural language processing model. a. Across areas that had high number of channels (≥ 25), the percentage of channels that were significantly correlated to the artificial network embeddings is plotted. The bar color denotes whether the percentage was calculated during speech production or comprehension, and the edge color labels the left or right hemisphere. Brain areas highlighted in bold represent those where the percentage of correlated channels significantly exceeded what would be expected by chance, which was focused on for the rest of the manuscript. **b**. Locations of channels that showed significant correlation to NLP embeddings during speech production *(top)* and comprehension *(bottom)*. **c**. The average of absolute R-values of correlated channels in natural conversation and control task when the participants were instructed to listen and repeat designed sentences. Gray bars label the R-value from all neural-NLP pairs. **d**. Plotting the percentage of correlated channels based on frequency bands for both language production and comprehension. **e**. Plotting the percentage of correlated channels based on the NLP layers from all areas in natural conversation *(left)*, and in control tasks with passive listening and repeating sentences *(right)*.

Overall, there were more channels with correlated activity from areas in the left hemisphere when compared to right (Chi-square test, statistic = 1214, p = 3.6 x 10^-266^, **Fig. S2a**). Further, across 17 brain areas with a high number of channels implanted (n ≥ 25, Method), there were 9 areas that showed significantly high percentage of channels (p < 0.01) that correlated to NLP embeddings, including areas in the temporal and frontal cortex, thalamus and limbic system structure (**Fig. 2a**). Thus, in subsequent analyses, we used these 9 areas as representative regions of interest. Specifically, and as might be expected, the highest ratio of channels with correlated activity was found in the left precentral cortex during speech production planning (21%) and left and right superior temporal cortex (23% and 25%, respectively) during comprehension (**Fig. S2a**). In addition, all frontal, temporal and mesial brain areas appeared to engage neural activity in multiple frequency bands for language encoding during both speaking and listening (**Fig. 2d, Fig. S2b**). These results and the general distribution of these correlated channels were largely consistent across participants (**Fig. S3**), regardless of their intelligence quotient (**Fig. S4**) and the length of the conversation (**Fig. S5**), and were also robust if assigned white matter locations to their nearest gray matter (**Fig. S6**). Together, these findings suggest that neural activity in diverse brain regions, especially frontotemporal networks, paralleled that of the NLP models, with similar neural representations of linguistic information captured by the models during speech production and comprehension.

### Generalization and robustness of the neural-NLP relations in natural conversation

To further ensure that the relationship between neural activity and those of the NLP model was language-related, we randomly permutated the neural activities over words to eliminate any linguistic information obtained by chance. When randomized, we found a significantly lower proportion of contacts that displayed selectivity (Chi-square test, statistic = 5002, p < 10^-100^, **Fig. S7a**) and a significant drop in the degree of correlation to R = 0.02.

Secondly, we show that these NLP-correlated activities cannot be explained by correlations to the low-level sound features. We reasoned that if the observed neural-NLP correlation originated from neural activities responding to low-level sound features, we would expect that the NLP-correlated channels exhibit a higher response to sound features such as the voice amplitude and pitch in comparison to the response of sound features from all channels (including those responses which do not correlate to NLP). However, the NLP-correlated channels responded similarly to both voice amplitude and pitch as compared to all recorded channels (2-side permutation test combined, amplitude: p = 0.41 for speak, p = 0.21 for listen; pitch: p = 0.79 for speak, p = 0.60 for listen; n = 10000).

Thirdly, to confirm that the above findings were generalizable, we compared the neural activity patterns to those of another NLP model trained on a different dataset. Here, we used a BERT (base) model with a bidirectional network architecture and trained on different linguistic materials *(41)*. Again, we found that the proportion of correlated channels was significantly higher than chance (Chi-square test, statistic = 15278, p < 10^-100^), and was greater than or equal to that observed with the GPT-2 model (**Fig. S7b**). Together, these findings suggest that the relationships between neural activity patterns in the brain and those of the language models are generalizable properties of neuronal responses.

Finally, we investigated the extent to which information uniquely associated with natural conversation differs from passively listening to sentences in block design tasks by instructing the participants to listen to sentences and repeat the words they heard. This task was designed to simulate the format of speaking and listening, but to replace the spontaneity of natural conversation with constrained materials (Methods). In total, we performed this task on 4 participants (465 channels) with each participant hearing 237 ± 6 (mean ± s.d) and speaking 233 ± 6 words on average. Next, we calculated the average of R-values (range 0 to 1) from the neural-NLP pairs that were significantly correlated in natural conversations. We found that the average correlation coefficients significantly decreased when the participants were passively involved in the pseudo conversation (0.13 to 0.09 for speech production, and 0.11 to 0.07 for comprehension, T-test, p < 2 x 10^-56^ for both, **Fig. 2c**). Therefore, these neural signatures were specifically correlated with conversation rather than, more simply, speaking and listening.

### Neural-NLP relations across frequency bands and NLP layers

The neural-NLP correlation not only spanned broadly across the frontal, temporal and mesial areas, but also occupied multiple frequency bands. Specifically, the percentage of correlated channels of each frequency band was consistently higher than chance for both comprehension and production (Chi-square test for each frequency, statistic > 13, p < 3 x 10^-4^; FDR corrected for multiple frequencies), with the highest percentage observed in middle gamma frequencies (70 - 110 Hz) for both language comprehension and production (11% and 14%, **Fig. 2d**). Further, these frequency patterns showed weak dependence on brain area for both production and comprehension (Repeat Measurement ANOVA, F(32) = 2.49, p = 0.063 for speak, F(32) = 3.3, p = 0.02 for listen, **Fig. S8**). For example, during speech production planning, 13% of channels in middle temporal cortex showed correlation to NLP embeddings in the high gamma band, while this only 5% showed similar relationships in the rostral middle frontal cortex. In addition, while many areas had a higher percentage of channels demonstrating correlations in middle gamma frequency during comprehension (e.g. 24% and 35% in middle and superior temporal cortex), other areas, including the hippocampus and amygdala, contained a higher percentage of channels with correlations in the alpha frequency (12% and 10% respectively). Together, these findings show that not only are the areas of activity widespread but the frequencies of neural activity that are relevant for language processing are also broad, suggesting parallel, multiscale neural dynamics that intracranial EEG is uniquely able to characterize compared to non-invasive fMRI method.

Neural-to-model correlations during conversation were also largely confined to distinct hidden layers within the language models. Whereas lower (input) network layers within the NLP model preferentially reflect information about individual words independent of their context, higher (output) layers reflect integrated compositional sentence-level information *(42-46)*. Therefore, to further examine the relationship between neural activity patterns and that of the NLP model, we calculated the percentage of channels correlating with each network layer, and found that neural activities preferentially aligned with higher network layers for both speech planning and comprehension (12% and 14% for the first and last 6 layers during speak, T-test statistic = -3.2, p = 1 x 10^-3^; 14% and 18% for comprehension, statistic = -4.3, p = 1.8 x 10^-5^, **Fig. 2e**). This neural distribution across layers was significantly different from those when the participants were passively listening versus articulating the sentences they heard (repeated measurement ANOVA, F(12) = 23.3, p = 4 x 10^-4^ for listening, F(12) = 5.6, p = 0.036 for speaking). Further, the specific layer distribution was dependent on the brain area (repeated measurement ANOVA, F(96) = 4.2, p < 1 x 10^-5^ for speak; F(96) = 6.7, p < 1 x 10^-5^ for listen, **Fig. S9**), with those from temporal lobe exhibiting higher layer-dependence compared to those in frontal cortex (temporal: 11.8% for first 6 layers, 15.0% for last 6 layers during production planning; frontal: 9.1%, 9.9%). In short, the majority of neural activity patterns likely reflect higher order contextual information rather than low level early processing or pre-speech motor planning.

### Neural activity patterns during speech production versus comprehension

While the above findings suggested a broad representation of linguistic information during production and comprehension, it remained unclear whether or how these representations overlapped spatially across the frontotemporal network. Therefore, to examine this in more detail, we tested the correlation between neural activity and the NLP models that were specific to either production or comprehension. Overall, we found a similar number of channels whose neural activities were correlated with that of the NLP models during speech production planning compared to comprehension (210 and 286 contacts, respectively; **Fig. 3a**) as well as a similar strength of correlation with the models (R = 0.12 vs. R = 0.11, **Fig. 3a**). Across the patients, there were 76 channels (18%) that showed shared responses for both speech production and comprehension, displaying some overlapping processing between the two activities. These overlapping representations were significantly higher than chance (Chi-square proportion test, statistics = 19, p = 1.3 x 10^-5^), although 82% of selective channels only responded during production or comprehension alone.

**Fig. 3.**
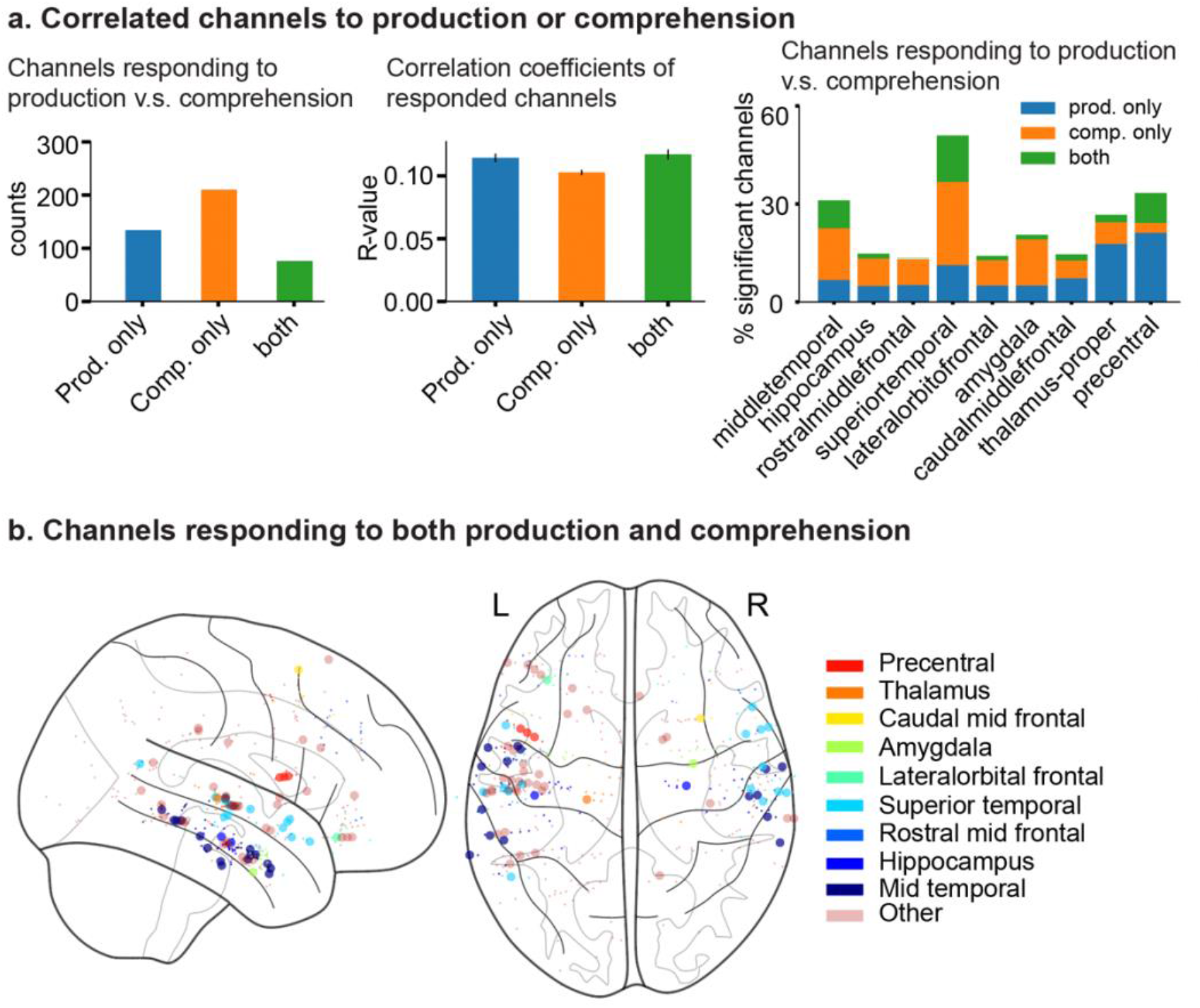
Comparing neural activities during language production and comprehension. **a**. Demonstration of channels selective to artificial embeddings during one or both production and comprehension *(left)*. Correlation coefficients are plotted by channels significantly correlated to artificial embeddings during comprehension or production *(middle)*. Percentage of correlated channels selective to language production and comprehension were plotted by their locations *(right)*. **b**. The locations of the channels that responded to both production and comprehension were shown in large dots, and the smaller dots showed the channels that only responded to one of comprehension or production.

Furthermore, different brain areas exhibited different extents of overlapping processes (**Fig. 3a, b**). For example, most channels selective to production were also selective to comprehension in the superior and middle temporal cortex (14% and 11% selective to both vs. production alone in superior temporal cortex; 8% and 7% in middle temporal cortex); Similarly, most channels selective to comprehension were selective to production in the precentral cortex (9% and 3% selective both and perception alone). In contrast, other areas, including rostral middle frontal cortex and lateral orbitofrontal cortex and amygdala, displayed minimal overlapping between speak and listen (≤ 1%). This inhomogeneous distribution of channels responding during either speech production, comprehension, or both cannot be attributed to randomness (Chi-square test considering the ratio of channels selective to both to the total number of channels, p = 2 x 10^-6^). Collectively, although the channels selective for speech production and comprehension exhibited certain degree of overlap across all examined brain areas, there was a significant inhomogeneity in their distribution, particularly within the specific areas we scrutinized.

### Neural activity during speaker-listener transitions correlated with that in response to the NLP model

A core aspect of conversation is the alternating transition between listening and speaking. Given the distinct neural processes underlying speech production and comprehension, we also examined activity during transitions between speaker and listener to ask whether these neural activity patterns aligned with specific transitions rather than established correlations for production or comprehension. To start, we tracked specific transition points at which the participants changed from listener-to-speaker or from speaker-to-listener during dialogue. These were the times when they specifically shifted between assimilating and communicating information (**Fig. 4a**) *(13)*. Across all brain areas, we found that 13% of channels displayed a significant change in activity during transitions from comprehension to production, whereas 12% displayed significant changes at the transitions to comprehension (**Fig. 4b**). Both proportions were significantly higher than random chance (Chi-square proportion test, p = 7 x 10^-88^ and p = 1 x 10^-64^), and most of these channels were only selective during natural conversation, compared to a control in which the participant was ‘forced’ to take turns (66% channels only selective to turn-taking to speak during natural conversations but not during designed turn-taking, whereas the percentage was 55% channels selective to natural turns to listen, **Fig. 4b**). In addition, the number of contacts at the transitions from production to comprehension was significantly smaller than the transition from comprehension to production at low frequency bands (alpha, beta bands, permutation test, p = 0.048, 0.035 respectively), whereas there were little differences at the mid- and high gamma bands (p = 0.39 and p = 1.0) between the two directions. These patterns were consistently found throughout the brain (**Fig. S10**). Further, the patterns also differed in polarity in terms of increases to decreases in activity. For example, there was a decrease in beta frequency activity in the superior temporal cortex during transition to comprehension, whereas channels from superior and middle temporal cortices demonstrated an increase in middle and high gamma bands (**Fig. S11**). Together, neural activities reflected transitions during turn-taking, showing diverse changes among brain areas and frequency bands.

**Fig. 4.**
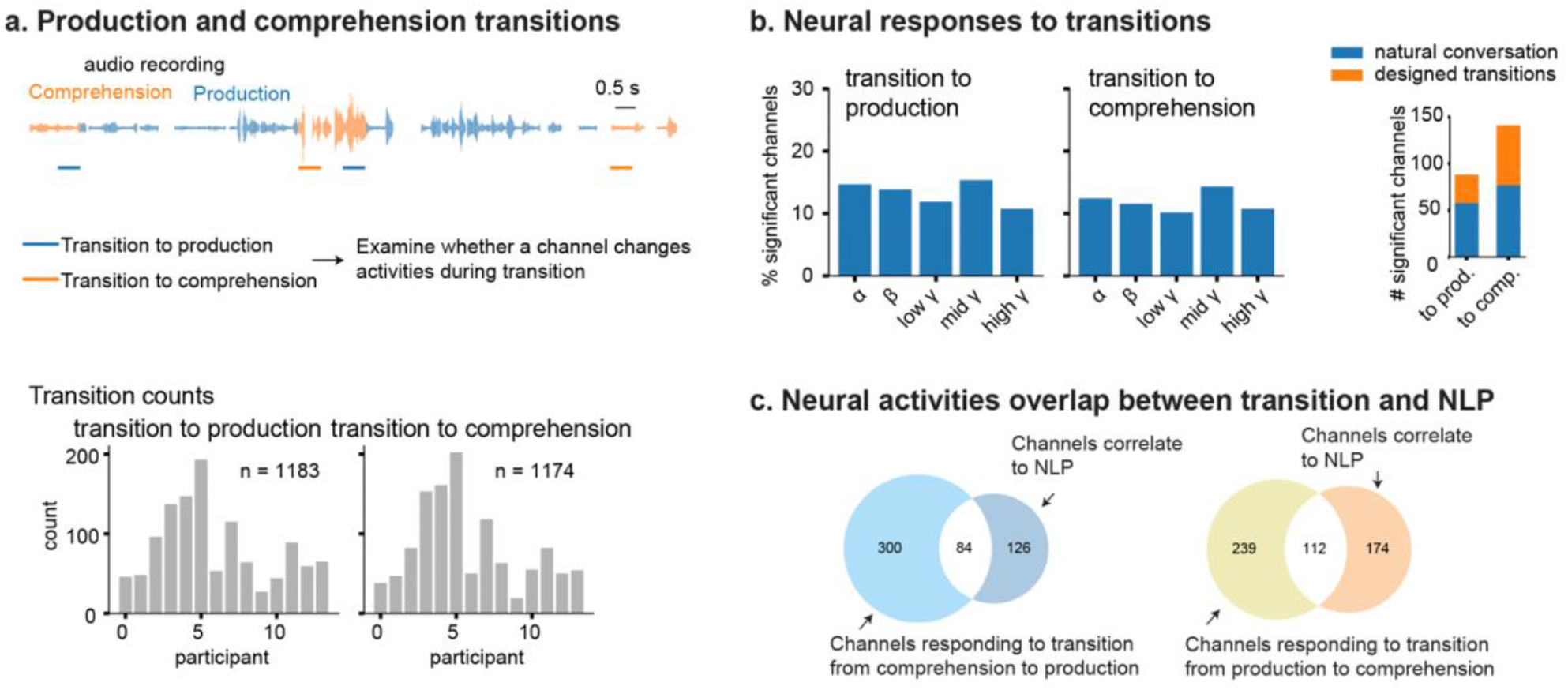
Neural activity changes with turn-takings during conversation. **a**. Audio waveform illustrates the transitions between production and comprehension *(top)* and the counts for turn number for each participant *(bottom)*. **b**. There are an average of 13% and 12% channels that showed significant changes of activities across all five bands during transitions, but transition to comprehension adopts a lower number of channels at low frequency bands (alpha, beta) compared to transition to production, whereas the mid- and high gamma bands are similar. The subplot on the *right* shows the channels that responded to turns during natural conversation compared to designed transitions. **c**. Comparing transition-responding channels to the ones that are significantly correlated to NLP during production, there is significant overlapping between NLP-response channels and the transition-response channels.

Finally, we asked whether neural responses to these speaker-listener transitions overlapped with those reflecting correlations with the NLP models. It is possible, for instance, that neural responses to these transitions used entirely separate subsets of neural patterns compared to those encoding the specific information being conveyed during conversation. We found, however, that 39% (112 out of 286) of the contacts whose activities correlated with that of the NLP models overlapped with those that displayed speaker-to-listener transitions (**Fig. 4c**), while 40% (84 out of 210) of the production-planning contacts displayed changes during listener-to-speaker transitions. This degree of overlap was markedly higher than that expected by chance (Chi-square contingency; p = 4.9 x 10^-14^ and 1.7 x 10^-22^ for each transition direction respectively). These neural activity patterns therefore appeared to reflect not only information of the conversations being held but also tracked the specific transitions between speakers at which times the ‘directionality’ of communication changed – elements that together would be necessary for our ability to converse.

## Discussion

Major challenges in understanding how our brain supports natural conversation include limitations in precisely recording the relevant neural activity, the difficulty of richly and accurately representing linguistic information, and the ability to place both of these problems in the context of natural human interactions *(47, 48)*. By comparing intracranially recorded neural patterns during natural dialogue between individuals to those of NLP models, we have taken a novel route toward overcoming these significant hurdles. We found that linguistic information as captured by the models could similarly explain neural activity patterns found in the brain when the participants were not only language comprehension, but also during language production. These neural patterns were carried across multiple frequency bands and showed heterogeneity over brain areas. These findings extended the previous demonstration of neural-artificial convergence focusing only on comprehension *(34, 49)* and revealed a shared neural language representation with speech production. In addition, the correlation between neural activity patterns and that of the NLP model’s, by contrast, were reduced during designed blocks of pseudo dialogue and largely absent when randomizing the neural data, together suggesting that these neural patterns reflected the specific linguistic information being conveyed in natural conversation.

The information being conveyed in speech production, though, did not simply mirror that of comprehension. Here, we found a higher ratio of channels involved in language processing in the temporal cortex during comprehension than production, whereas the precentral cortex was biased towards production planning, suggesting distributed processing with varying functional tendencies for different brain regions. In addition, most neural activities throughout frontal and temporal lobes reflected either production or comprehension, with less than 20% of single channels involved in both. Together, these findings suggest the presence of a core set of locations that respond selectively to both language modalities, in addition to areas reflecting exclusively one of the processes. Although previous fMRI work has suggested partially shared brain areas underlying language comprehension and production *(50-53)*, sEEG recordings offer direct measurement of neural activities that have high temporal resolution and can be applied, leveraging deep learning approaches, to natural conversation. This provides a richer and more complete view of the neural basis of normal human conversational than has been achieved previously.

Further, when tracking neural responses during comprehension-production transitions, we observed that many brain areas significantly changed their activity during turn-taking; a process surprisingly similar to that observed in prior animal and human models of communication *(13, 54)*. More notably, these neural changes closely overlapped with brain areas that displayed linguistic information as identified by neural-to-model correlations, together suggesting that these response patterns reflected the process of communication rather than simply the act of listening or speaking.

A final striking finding was the relationship between neural activities and the activities of specific nodes in the NLP models *(34, 49, 55)*. Overall, we found that the neural dynamics of most areas preferably correlated with representations in the middle and higher layers of the NLP model, suggesting that these activity patterns reflected contextual or sentence-level information integration over the course of conversation rather than simply related to individual words which would be represented by lower layers. Notably, these layer distributions were significantly altered when the participants passively listening and repeating sentences, indicating differences on processing mechanics during block designed language tasks compared to more natural language.

Taken together, using a combination of intracranial recordings, NLP models, and naturalistic conversation, our findings reveal a set of collective neural processes that support conversation in humans and a detailed organization of neural patterns that allow linguistic information to be shared across speakers. These observations provide a prospective neural framework for understanding the detailed neuronal computation underlying verbal communication in humans.

## Data availability

Data will be made publicly available on public repositories.

## Code availability

Codes will be made publicly available on GitHub.

## Acknowledgements

J.C is supported by the American Association of University Women, Z.M.W. is supported by NIH R01DC019653 and NIH U01NS123130 and S.S.C., A.E.H., A.C.P., D.J.S. are supported by NIH U01NS098968.

## Author contributions

A.E.H., J.C., Z.M.W. and S.S.C. designed the study; J.C., Z.M.W. and S.S.C. conceived and designed the language processing analysis platform and its implementation; A.E.H., D.J.S., A.C.P., J.C. and S.S.C. collected data; J.C., A.E.H., A.C.P., T.X. and S.S.C. pre-processed data; J.C., Z.M.W. and S.S.C. performed analysis and modeling and prepared the manuscript. All authors edited the manuscript.

