## Supplementary Material for "Natural language processing models reveal neural dynamics of human conversation"

### MATERIALS AND METHODS

In this study, all data acquisition and analyses were approved by the Partners Human Research Committee Institutional Review Board (currently Massachusetts General Brigham Institutional Review Board). Before their enrolment in the study, participants were informed that their participation would not impact their clinical treatment, and that they could withdraw their participation at any time without impact of their clinical care. Consent was obtained from all study participants.

##### Stereo EEG recording

We recorded neurophysiological data from fourteen intractable epilepsy participants when they went through invasive monitoring for localization of the brain areas that onset their seizures (**Table S1**). A team of clinicians who were independent of this study determined the decision to implant electrodes, as well as the types of electrodes and the areas for implantation solely based on clinical grounds (**Table S2**). We recorded neuronal activities using one or two neural signal processor systems (128 to 256 channels, NSP, Cerebus, Blackrock Microsystems). These raw voltages were digitized at a 2 kHz resolution and then filtered online to capture local field potentials (low-pass filter, 1 kHz cutoff). If two systems were used for recording, we aligned them within a temporal error of 10 ms.

##### Natural conversation

We recorded the speech perception and production of the participants from conversations between the participant and the experimenter (> 15 minutes). These conversations varied broadly in topic and theme to allow the participants to actively engage in dialogue whereby they both listened and spoke (**Table S3**). For example, conversations would involve real-world topics such as a movie that was recently watched or interpersonal thoughts and events shared between the experimenter and the subject.

##### Control task with passive listening and speaking sentence

To investigate to which extent information is uniquely associated with natural conversation compared to passively listening and speaking sentences in block design tasks, we performed the following control experiment: Participants were instructed to listen to sentences and passively repeat the words they heard. This task was designed to simulate the format of a natural conversation, but replacing the content of the dialogue into designed materials. In total, this trial contains 240 words with 8 words per sentence.

##### Speech recordings

We used an audio recorder (DR-40X by TASCAM) to record the conversation. The audios were synchronized with the intracranial activity at a millisecond resolution by adding an analog input channel to the NSP (Blackrock Microsystems). Audio recordings for each participant were firstly automatically transcribed using whisperX (*1*), then manually adjusted to align the time stamp for each word and manually assigned the speaker identity (participant or experimenter) using either Audacity (The Audacity Team, version 3) or SpeechScribe, a Python application developed in house for manual speech annotation. Any words or sentences containing participants' personal information were removed and replaced by a set of other names and digits.

#### Estimation of number of channels

To estimate sample size (number of channels) to achieve 80% statistical power, we used the test for one proportion (*2*). Specifically, we followed the equation below to estimate the sample size $n$ based on the probability that a channel is significant $p_1$, chance level $p_0 = 0.05$, z-value for the threshold $\alpha = 0.05$ and the power $\beta = 0.8$:

$$n = \left( \frac{z_{\alpha/2}\sqrt{p_0(1-p_0)} + z_\beta\sqrt{p_1(1-p_1)}}{p_1 - p_0} \right)$$

We estimated $p_1$ with a range between 10% to 40%. The estimated sample size increased from 6 for $p_1 = 0.4$ to 185 for $p_1 = 0.1$. Therefore, the recorded number of channels after bipolar referencing, 1910, is much higher than the sample size required for 80% power.

#### Electrode localization

We adopted a combined surface registration and volumetric process to identify each electrode's anatomical location within the brain (*3*, *4*). We first used FreeSurfer (*5*) to align preoperative MRI data with a postoperative CT scan, then we manually transformed electrode locations identified from the CT scan into the MRI space (*6*). We mapped each electrode to a set of brain regions defined in the DKT atlas using an electrode labeling algorithm (ELA) (*7-10*). The electrodes were assigned with an additionally probability of being situated in white matter (**Table S2**). Electrode contacts were then mapped to MNI space using Fieldtrip volumetric morphing tools (*11*).

#### Bipolar referencing

Neural data re-referenced into a 'bipolar' configuration by taking the voltage differences between adjacent channels from the same electrode array in MATLAB (MathWorks). The importing of neural data was assisted by the NPMK toolbox (Blackrock Microsystems), and then processed in Matlab (6 participants) or in Python (8 participants). Channels that showed no variance or regular 60 Hz oscillation across the whole recording period were excluded from further analysis. The voltages were further decimated to 1 kHz. In total, we obtained 2234 electrode pairs from fourteen participants.

#### Interictal epileptiform discharge detection

Though no seizure occurred during the language tasks, interictal discharges may be common. In order to remove these possible large amplitudes confounds from further data analysis, we detected epileptiform discharges automatically (*12*) and removed channels in which there were a high rate of epileptiform events (> 6.5 discharges/min). This resulted in a total of 1910 bipolar channels remaining from fourteen participants.

#### Frequency bands computation and alignment to words

LFPs were further processed for amplitudes at alpha (8-13 Hz), beta (13-30), low gamma (30-55), mid-gamma (70-110), high gamma (130-170) frequencies using the scipy package for signal processing in Python with the following steps (**Fig. S1**): To construct the envelope of each frequency band, the voltages were filtered by a Chebyshev type II filter with $4^{th}$ order and 40 dB attenuation for each frequency band, and a Hilbert transformation was further applied. Next, outliers of the envelopes were capped to $q3 + c \cdot (q3 - q1)$ and floored to $q1 - c \cdot (q3 - q1)$, where *q1* and *q3* are the first and third quartiles, and *c* = 5 determines the threshold of outliers. To compute the neuronal activities for each word, envelopes were initially aligned to the auditory signals. For each word, the neuronal activities at each frequency band were computed as the average of the envelope over a 0.5 second window before word onset for speech production planning and after word onset for comprehension. After this step, for each frequency band of a channel, we obtained an array of neural activities at the same dimension as the word number.

**De-autocorrelation**

EEG signal is known to possess high long-range temporal correlations and 1/f-like power distribution (*13, 14*). In our study, neural activities by words indeed showed strong autocorrelations at all frequency bands we examined. After we aligned the envelop for each frequency to the onset of words, and treated the progressing of words as a time series, we found that the first order autocorrelation (AR1) dominated, with an average across all channels for each frequency band higher than 0.4 during speak and listen. These autocorrelations might result in spurious high correlation to NLP embedding if we had directly calculated the correlation. Therefore, we used a standard auto-correlation estimation called Cochrane-Orcutt method to process AR(1) in the neural activity (*15*). To avoid heavy computation, we estimated $\rho$ as the AR(1) term from the neural activities and the NLP embeddings respectively. Specifically, the first order autocorrelation of neural activities as the sequence of words was calculated for each frequency band:

$$\rho = \mathrm{corr}(X_i, X_{i-1}), \quad (1)$$

where $X_i$ is the neuronal activity for the *i*-th word. Then we removed the AR(1) by transforming the envelopes by

$$\tilde{X}_i = X_i - \rho X_{i-1}. \quad (2)$$

Next, we further treated the outliers with *c* = 1.5 using the same method as described in the previous paragraph to ensure that our findings were not driven by just a few points. In this way, we only used one variable $\rho$, and it is only counted the degree of freedom of 1. This autocorrelation does not directly reflect any information specific to language (example, part of speech, lexical semantics, etc).

**Natural language processing (NLP) network embeddings**

We selected a Generative Pre-trained Transformer (GPT-2, small) model to encode language information and compare to brain activities (*16, 17*): The OpenAI GPT-2 model has been applied in previous work to multiple brain imaging studies and has been shown capable of capturing variance of brain activities from multiple areas during language comprehension (*18-20*). This model was trained on a combination of text datasets that were built on a wide range of natural language demonstrations including diverse domains and contexts, hence, the GPT-2 model was able to process natural dialogue with informal colloquial terms (e.g., “gonna” rather than “going to”) often encountered during real-world conversations (*21-24*). We directly loaded the tokenizer (mapping of a word to a vector) and the model from the pre-trained GPT-2 small model in PyTorch via huggingface without fine-tuning (*25*). The model was built on 12 structurally identical modules in sequence and input sentences were represented between layers by hidden embeddings, which were 768-dimension vectors for each word per layer (*i.e.* 768 nodes or artificial neurons). The model outputs 13 sets of hidden embeddings with 768 dimensions for each word of a sentence. These 13 embeddings include the embeddings ‘output’ of the 12$^{th}$

hidden layers and the ‘input’ embedding to the first module. To capture the contextual information of the discourse for each word, we input words with a moving window of 600 words (concatenation of current and previous words) with stride of 10 words to NLP model to get the embeddings. For words corresponding to more than one tokens (part of words), the embeddings were obtained from the last token of the word. After this step, we got the embeddings with the dimension of number of words x 768 nodes x 13 layers for each participant.

**Correlating neural activities to NLP model embeddings**
To examine how brain activities involve in language processing, we correlated neural activities to artificial embeddings by checking all neural-artificial pairs. Specifically, after de-auto-correlating the artificial embeddings following the same procedures described in Eq. (2) and treated outliers using the method described above with $c = 1.5$, we fit the neural activities (as a function of words) per frequency band of an channel using the artificial embeddings (on the same words) of a node to a linear regression, and obtained the Pearson correlation and the p-value of whether they were significantly corelated ($H_0$: slope = 0). Any neural-artificial pair with p-values lower than 0.05 (Bonferroni corrected for multiple comparison over 768 dimensions) was considered as correlated. Then for a given brain area, we examined whether there were significant number of channels correlated to the NLP model based on Chi-square test ($p < 2.5 \times 10^{-3}$, Bonferroni corrected for 13 layers).

**Grouping brain areas based on hemispheres and participants handedness**
Given that some participants in this study were left-handed, we further examined the neural responses to language based on the hemispheres from participants of different handedness. Interestingly, among the left-handed subjects, correlated channels from left-hemisphere were still significantly higher than those from the right hemisphere (**Fig. S2b**, 2-side permutation test with combining frequency bands and layers, $p < 10^{-4}$). Hence, instead of grouping hemispheres based on the handedness, we combined all participants and separated channels by left and right hemisphere regardless of their handedness in **Fig. 2**.

**Control with BERT (base) model as another NLP model**
To ensure the observed neural-artificial correlation was generalizable across embeddings from different NLP models, we calculated the neural correlation to a pre-trained BERT (base) model (*26*). BERT (base) model was composed of 12 layers with similar architecture (transformer modules) to the GPT-2 model and was independently trained on different language corpus. The modules in BERT model examined pairs of words including both past and future words. Due to this bi-directional pairing design, we separated language materials by sentences and obtained the embeddings of the words by inputting one sentence at a time. All the other procedures were exactly the same as that of the GPT-2 model. Note that the percentage of BERT-correlated channels after words articulated was higher than using GPT-2 model (**Fig. S7b**). This was possibly because the BERT model contained future word information from the same sentence, so the high percentage may come from articulation planning of future words in the sentence.

**Other controls of robustness of the neural-NLP correlation**
To further ensure that the observed neural-NLP correlations were robust and not driven by one participant or the length of the conversation, and they were not dependent on the intelligence of participants, we performed the following analysis. First, to ensure that the observed neural

responses were not driven by one participant, we performed the same analysis, but removed one participant each iteration of the analysis, and examined if there was any significant change of the percentage of responding channels. The result is shown in **Fig. S3**. A one-way ANOVA was applied to examine whether removing a patient caused a significant change of the ratio. No removal of a participant resulted in significant change of the correlated channel ratio. Second, we showed that the number of words that a participant spoke and listened to was not correlated with the percentage of responding channels. Specifically, we sorted participants based on the total number of words they speak and listen, and plotted the percentage of responding channels in **Fig. S3**. There is no correlation with the percentage of channels to the rank of words by participants ($p = 0.95$). This suggests that the sample size is not a factor contributing to our results, hence having a longer conversation is not expected to have an impact on the percentage of the responding channels. Finally, we performed full scale intelligence quotient (IQ) tests on 9 out of 14 participants. Based on this, we examined the percentage of channels that were significantly correlated to NLP embeddings for each of these participants, and performed a linear regression to test whether there is any significant correlation between the neural activities and the IQs. In **Fig. S4**, each pair of bars represent a participant, and the location of these bars are plotted based on their IQ values. There is no significant correlation between the percentage of channels and participants IQ ($p = 0.10$ for speak, and $p = 0.24$ for listen). Therefore, based on the number of participants we have, the percentage of responding channels seemed not impacted by their cognitive ability.

**Neural activities correlate to turn-taking**

We investigated the turn-taking properties of a conversation by examining whether any channels show significant changes in each frequency envelope during speaker-listener transitions. For transitions from comprehension to articulation, we averaged neural activities at each frequency band using a 0.5 s time window before the onset of the first word articulation; For transitions to comprehension, we averaged neural activities from 0.5 s window after the onset of the first word perception. The average activities at each frequency band were compared to the activities during established comprehension using a T-test from a Scipy package (ttest_ind) and the threshold of p-value to determine whether the activities changed significantly was set to be 0.05. Because speaker-listener transitions had much longer intervals compared to a word being articulated or perceived, we did not perform de-autocorrelation for this analysis. Similar to the method in the previous section, we selected the band that showed the lowest p-values and attributed each channel responding to transitions with the selected frequency band.

To examine whether the number of channels was significantly different from speaker-listener transition to listener-speaker transition, we used two neural populations with each including all channels and we labelled each channel by whether or not it was significantly responding to one direction of the transitions or the other. Next, we used these populations to perform a 2-side permutation test: We concatenated the two populations and randomly sampled from the mixture for a given number of times, with each sampling contained the same number of observations as the original two populations. Then for each sampling, the absolute difference between the average values of the two sets of random sampling was calculated, and this value was compared to the absolute difference from the actual populations. After repeating these steps of sampling, the p-value was then defined as the number of times when the difference of random sampling from mixed population was greater than that from the actual population, divided by the number of drawings.

### Supplementary Tables

**Table S1.** Summary of study participants, including sex and handedness.

| Subject | Handed-ness | Sex |
|---|---|---|
| Participant 1 | right | female |
| Participant 2 | right | male |
| Participant 3 | right | female |
| Participant 4 | right | male |
| Participant 5 | right | male |
| Participant 6 | left | female |
| Participant 7 | left | male |
| Participant 8 | left | male |
| Participant 9 | right | female |
| Participant 10 | right | female |
| Participant 11 | left | female |
| Participant 12 | right | male |
| Participant 13 | right | male |
| Participant 14 | right | male |

**Table S2.** Channel distribution across brain areas

| area | counts |
|---|---|
| **Cerebral-white-matter** | 576 |
| **Middle temporal** | 177 |
| **Hippocampus** | 142 |
| **Rostral middle frontal** | 115 |
| **Superior temporal** | 106 |
| **Amygdala** | 78 |
| **Lateral orbitofrontal** | 78 |
| **Caudal middle frontal** | 55 |
| **Putamen** | 46 |
| **Thalamus** | 45 |
| **Inferior temporal** | 37 |
| **Precentral** | 33 |
| **Pars triangularis** | 33 |
| **Caudate** | 31 |
| **Insula** | 29 |
| **Inferior parietal** | 28 |
| **Supramarginal** | 26 |
| **Superior frontal** | 26 |
| **Caudal anterior cingulate** | 24 |
| **Parahippocampal** | 23 |
| **Fusiform** | 22 |
| **Pars orbitalis** | 20 |
| **Medial orbitofrontal** | 20 |
| **Precuneus** | 19 |
| **Isthmus cingulate** | 17 |
| **Lingual** | 16 |
| **Superior parietal** | 14 |
| **Pallidum** | 12 |
| **Lateral occipital** | 12 |
| **Rostral anterior cingulate** | 9 |
| **Transverse temporal** | 7 |
| **Postcentral** | 6 |
| **Accumbens-area** | 6 |
| **Posterior cingulate** | 5 |
| **Pericalcarine** | 5 |
| **Cerebellum-cortex** | 5 |

| | |
|---|---|
| **Entorhinal** | 4 |
| **Parsopercularis** | 2 |
| **Cuneus** | 1 |

**Table S3a**. Example 1 of the natural dialogue between experimenter (E) and participant (P).

| Speaker | Sentences |
|---|---|
| E | Okay Batman would make a great dad |
| P | I just saw the new Batman movie the Lego Batman movie |
| E | Any good |
| P | Yeah it's pretty good |
| E | I think like the thing has there been one before |

**Table S3b**. Example 2 of the natural dialogue between experimenter (E) and participant (P).

| Speaker | Sentences |
|---|---|
| E | I was gonna ask when's the last time a joke made you laugh but technically |
| P | Oh I laugh all the time at jokes |
| E | Okay |
| P | Even the dumb ones honestly |
| E | Do you have a favorite comedian or |
| P | I think ah not really I like Seinfeld a lot |
| E | Okay |
| P | this one guy uh Louis CK |
| E | Louis CK is awesome |
| P | He's pretty funny |
| E | I think he actually comes from Boston |
| P | Yeah he he comes from around Boston yeah |

**Table S3c**. Example 3 of the natural dialogue between experimenter (E) and participant (P).

| Speaker | Sentences |
|---|---|
| P | I watch like Top Chef uh a few times and like Hell's kitchen and stuff |
| E | What's Top Chef |
| P | Those are pretty good |
| E | So |
| P | It's fun where like they get a bunch of people together and they give them a bunch of like challenges and then they vote them off one by one |
| E | No okay it's like survivor style |
| P | Yeah basically |

### Supplementary Figures

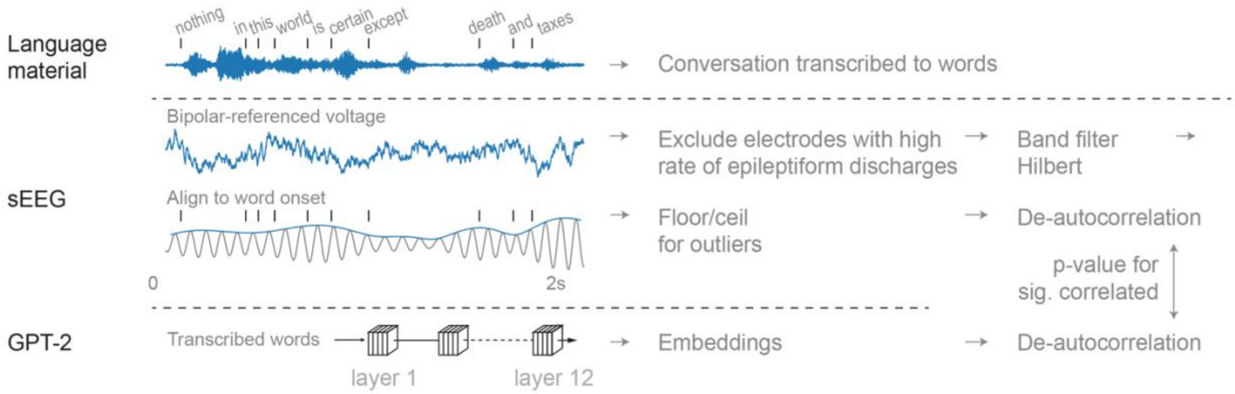


**Figure S1. a. Illustration of neural activity pre-processing and comparison to the GPT-2 model.** Audios of language materials were simultaneously recorded with all neural channels and were transcribed to words. Bipolar-referenced LFPs were processed for envelops at the frequency bands of alpha (8-13 Hz), beta (13-30), low gamma (30-55), mid-gamma (70-110), high gamma (130-170). Meanwhile, the transcribed words were input to a pre-trained GPT-2 model. We quantified the degree to which the artificial activities of specific nodes in the language model were predictive of neural activities obtained through depth electrode recordings by examining the correlation between the de-autocorrelated embeddings and the de-autocorrelated LFPs at each frequency band.

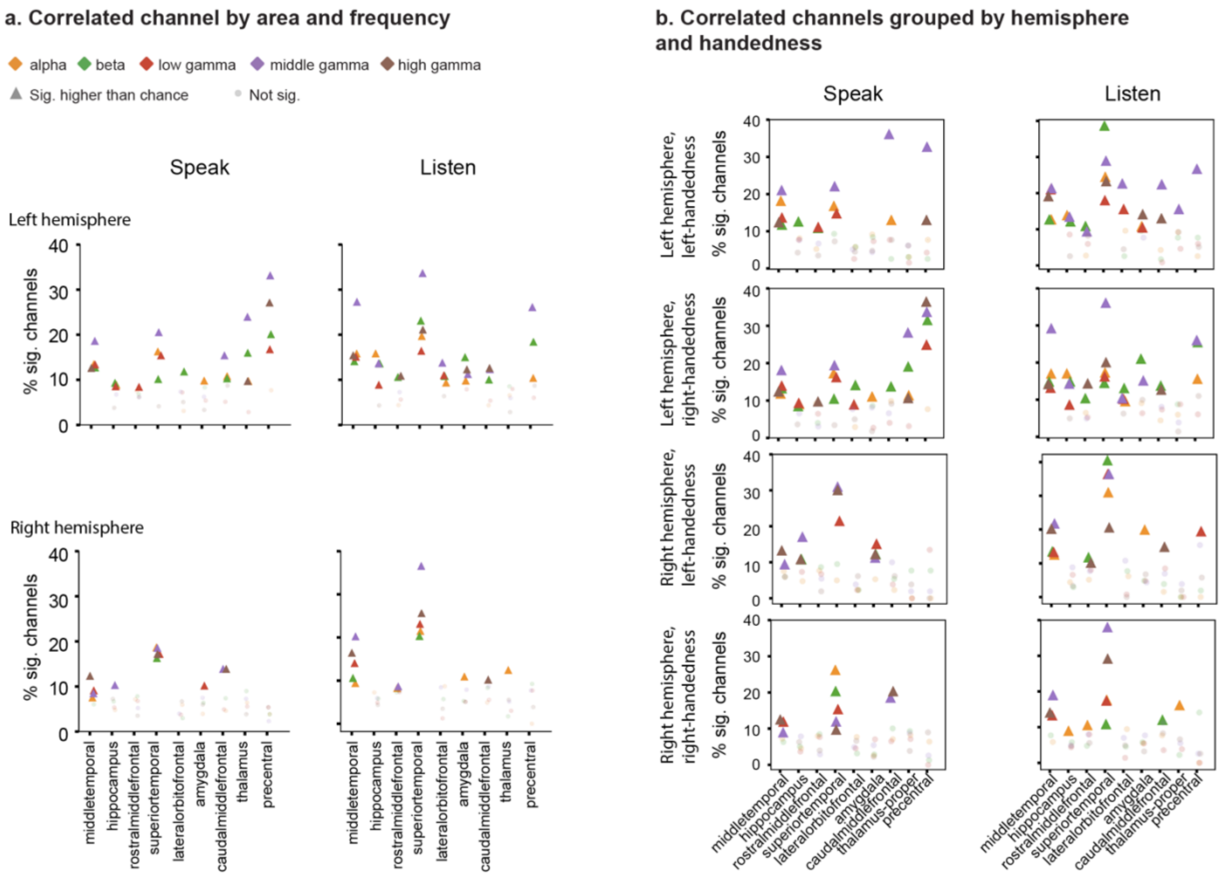


**Figure S2. Artificial-neural activity correlations by brain areas. a.** The percentage of channels with significant correlation to NLP model were illustrated by frequency bands, hemispheres, and brain areas. **b.** These channels were further grouped based on the handedness of the participants. Among left-handed participants, there were higher percentage of correlated channels in left hemisphere compared to the right (permutation test, $p < 10^{-4}$).

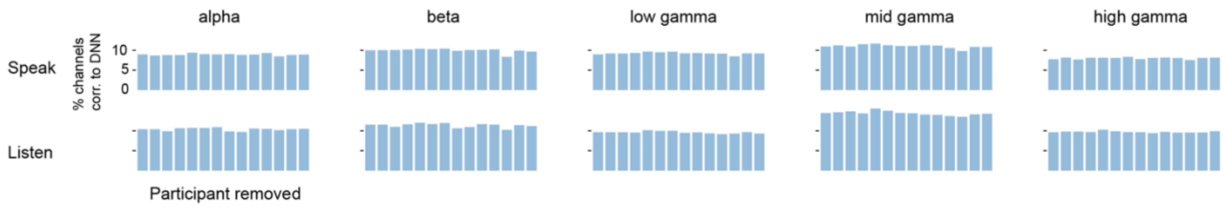


**Figure S3. Percentage of channels correlating to NLP model embeddings by removing a participant.** To ensure the ratio of channels significantly correlated to NLP was not caused by a single participant, we removed each participant (x-axis) and plot the percentage for both production (top row) and perception (bottom row) at different frequency bands. An one-way ANOVA was applied to examine whether removing a patient caused a significant change of the ratio, and all p-values were above 0.75.

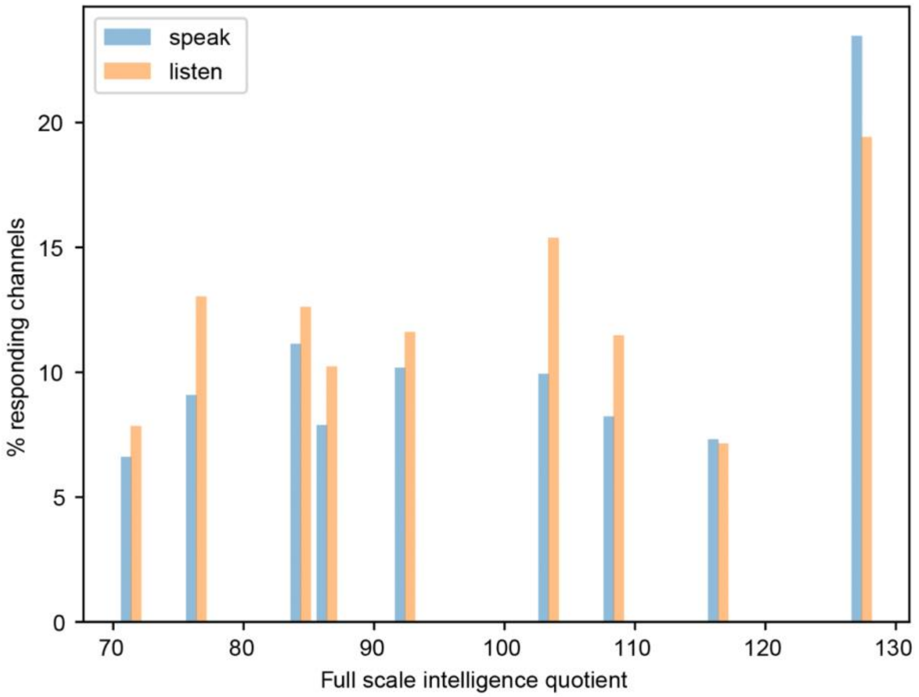


**Figure S4. Responding channel percentage versus participant's intelligence quotient (IQ).** To examine whether the neural response is correlated to the IQ score for the participants, we performed full scale intelligence quotient test for 9 out of 14 participants, and calculated the percentage of responding channels as a function of their IQ. Each pair of bars represent a participant, and the location of these bars are plotted based on their IQ values. There is no significant correlation between the percentage of channels and participants IQ ($p = 0.10$ for speak, and $p = 0.24$ for listen).

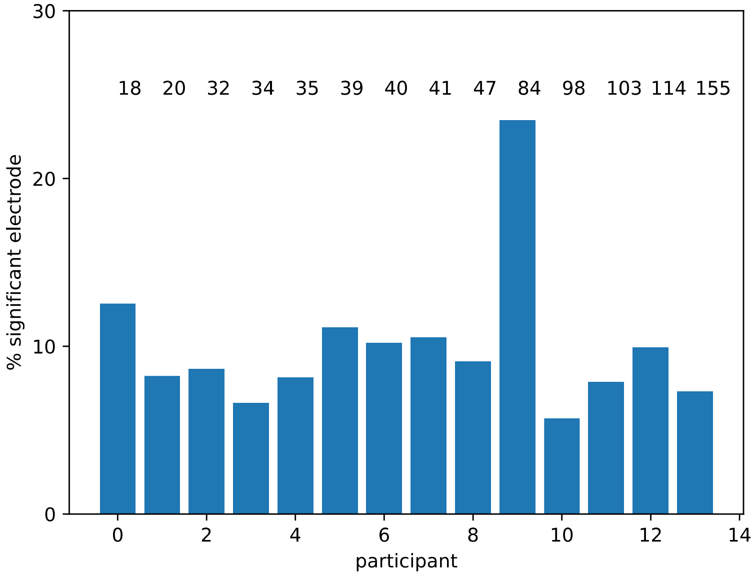


**Figure S5. Percentage of significant channels is not impacted by the number of words.** Participants were sorted based on the total number of words they speak and listen, and the percentage of responding channels are plotted. The numbers on top of each bar indicate the total number of words (x 100). There is no correlation with the percentage of channels to the rank of participants (p = 0.95). This suggests that the sample size is not a factor contributing to our results, hence having a longer conversation does not seem to have an impact on the percentage of the responding channels.

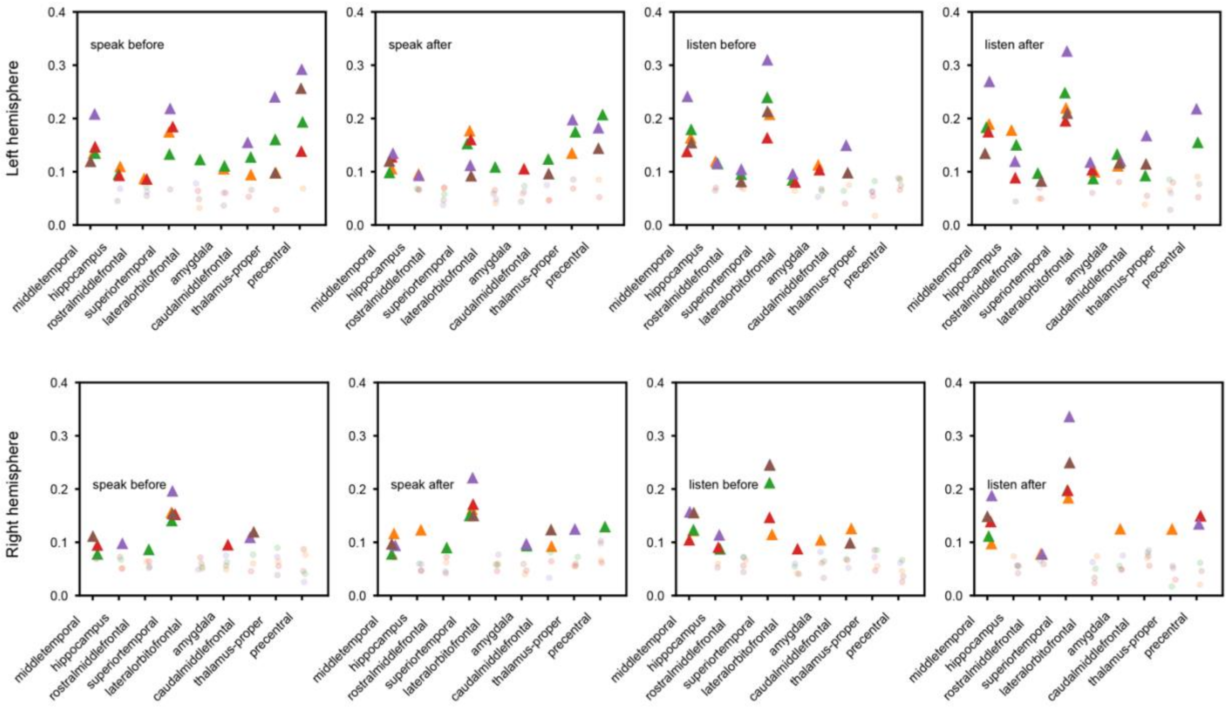


**Figure S6. Controls for the categorization of white matter.** In the main analysis, brain areas identified as white matter were categorized as 'white matter'. Since nearby gray matters contribute more to EEG, another way to categorize brain areas is to assign white matter locations based on the nearest gray matter. Here we show that the new categories without white matter result very similar ratios of channels that were significantly correlated to the deep learning model compared to excluding these channels.

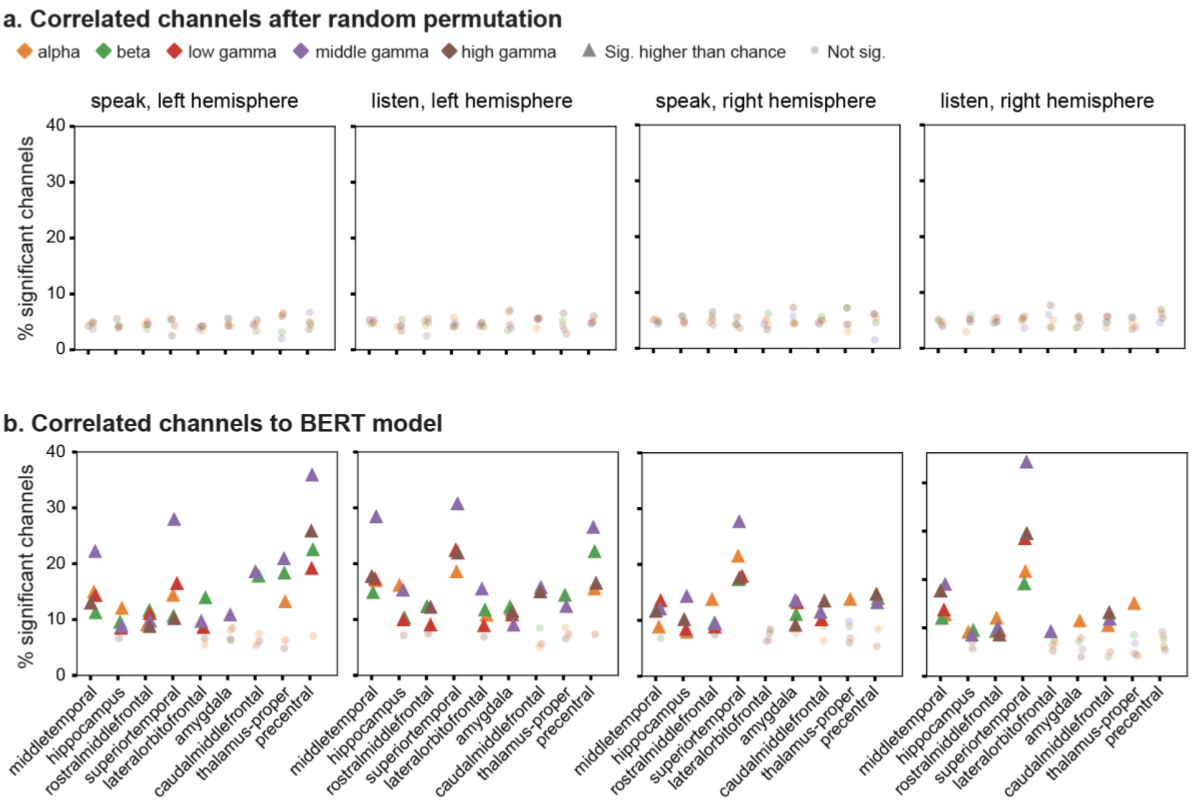


**Figure S7. Controls for robustness and generalization. a.** A control with the permutated NLP embeddings showed that correlated channels were homogeneously distributed at chance-level (5%), which were significantly lower than those without permutation (Chi-square test, statistic = 15278, $p < 10^{-100}$). **b.** Percentage of correlated channels to another pre-trained NLP model (BERT) showed similar or higher percentage to the pre-trained GPT-2 model, suggesting the neural-artificial convergence is generalizable.

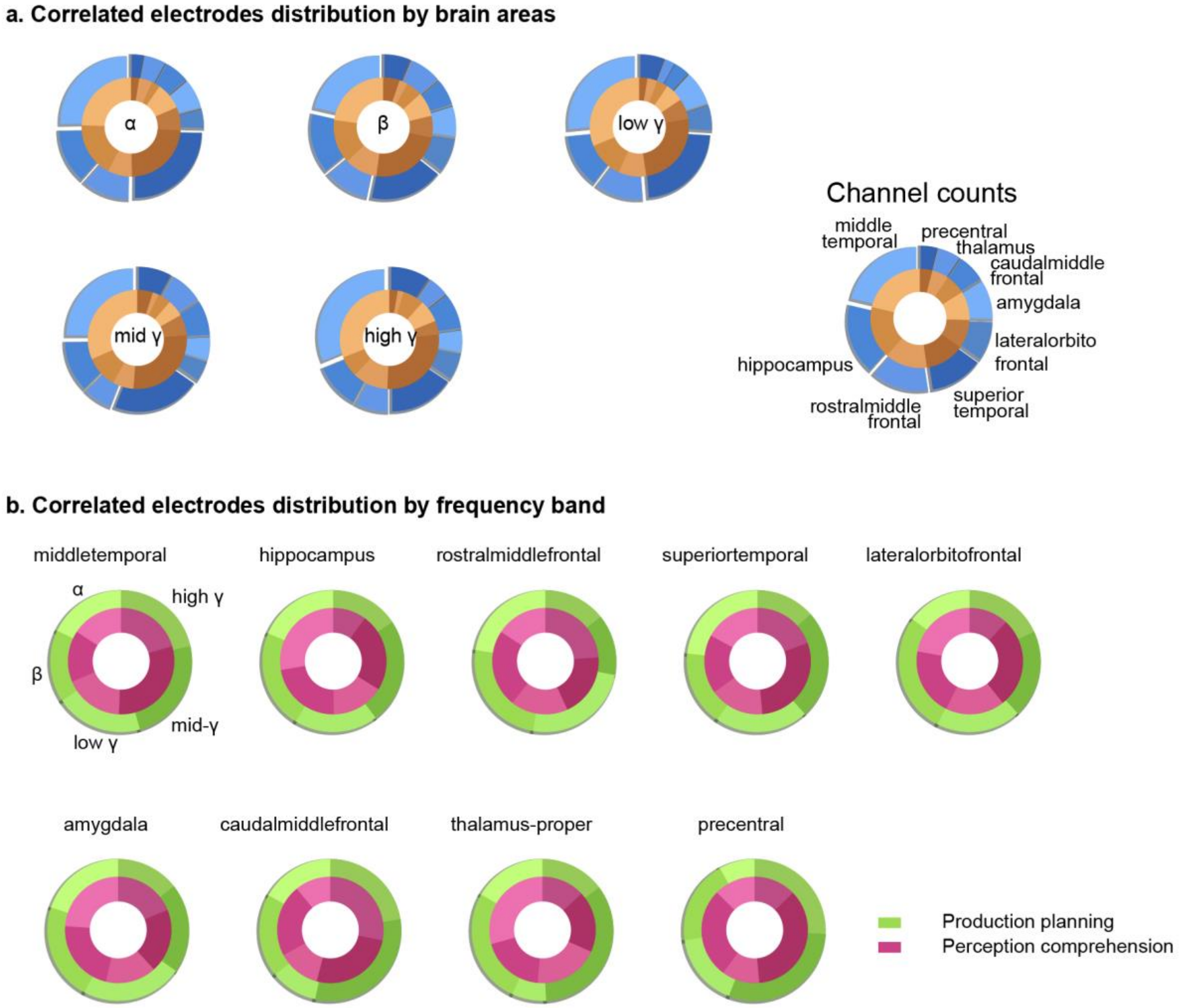


**Figure S8. NLP-correlated channels across brain areas and frequency bands. a.** Channels that were significantly correlated to NLP were grouped by brain areas and plotted across different frequency bands. **b.** Channels that were significantly correlated to NLP were grouped by frequency bands and plotted by different brain areas.

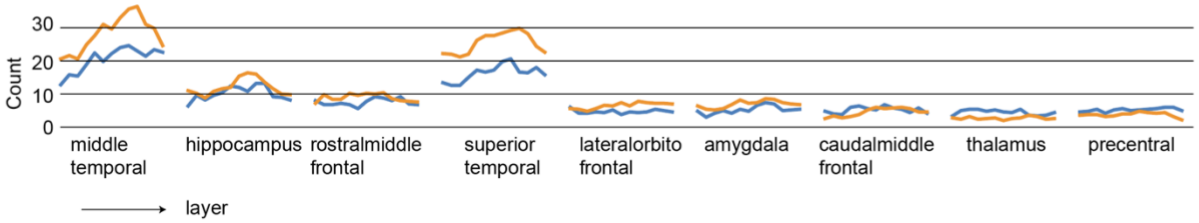


**Figure S9. Correlated channels across various NLP layers grouped by brain areas.** Channels correlated to NLP were plotted based on the NLP layers from various areas in natural conversation.

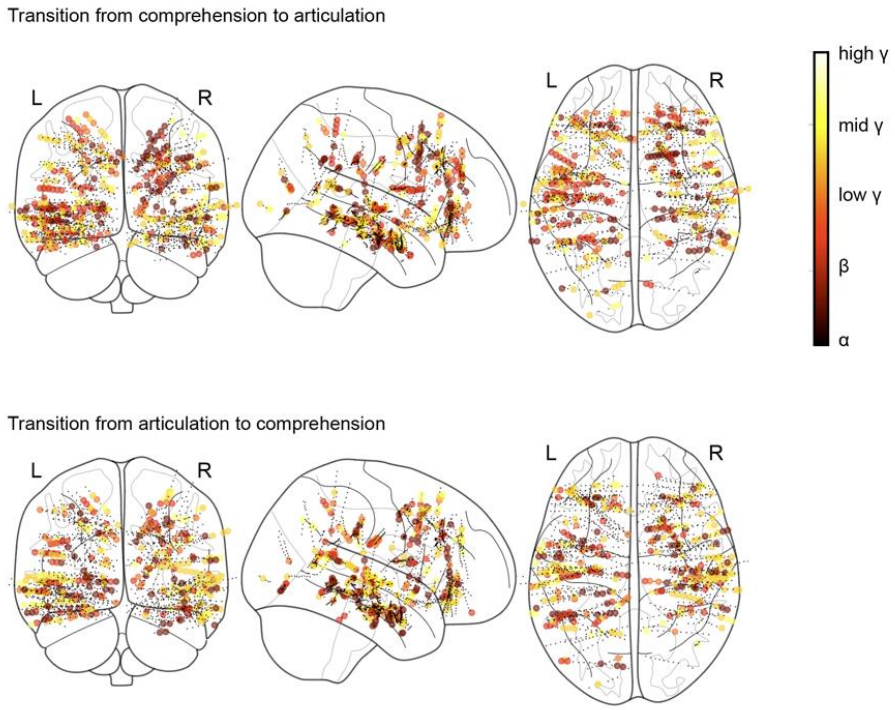


**Figure S10. Channels with frequency bands significantly changed during speaker-listener transitions.** Mapping of channels by their frequency bands that showed most significant changes during transitions.

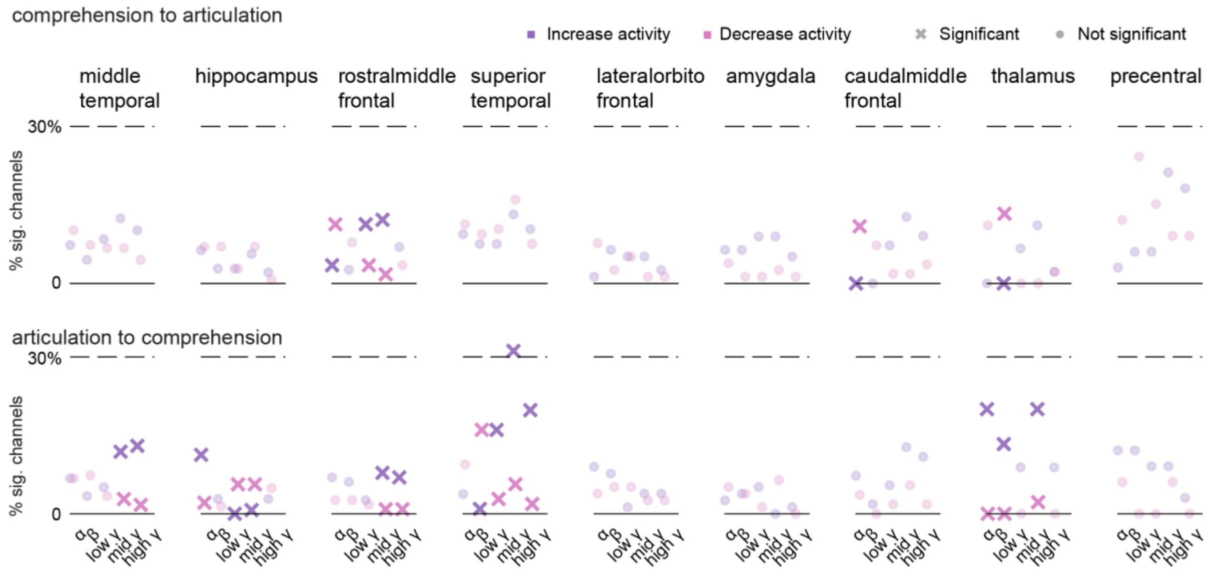


**Figure S11. Neural activity changes with turn-takings during conversation.** Percentage of channels that show increased activity (purple) and decreased activity (pink) by different the brain areas and the frequency bands.
